# Partitioning biota along the Elbe River estuary: where are the community transitions?

**DOI:** 10.1101/2024.05.13.593883

**Authors:** Benjamin B. Branoff, Luise Grüterich, Monica Wilson, Sven Patrik Tobias-Hunefeldt, Youssef Saadaoui, Julian Mittmann-Goetsch, Friederike Neiske, Fay Lexmond, Joscha N. Becker, Hans-Peter Grossart, Philipp Porada, Wolfgang R. Streit, Annette Eschenbach, Lars Kutzbach, Kai Jensen

**Affiliations:** Universität Hamburg; Faculty of Mathematics, Informatics and Natural Sciences; Department Biology; Institute of Plant Science and Microbiology; Ohnhorststr. 18 22609 Hamburg, Germany; Leibniz-Institute of Freshwater Ecology and Inland Fisheries (IGB); Experimental Limnology; Zur Alten Fischerhütte 2 16775 Stechlin OT: Neuglobsow, Germany; Universität Hamburg; Institute of Soil Science, Allende-Platz 2 20146 Hamburg, Germany

**Keywords:** Salinity, Community Assemblages, Ecotone, Brackish

## Abstract

Estuarine zonation and community assemblages are frequently correlated with salinity, although the extent or nature of this correlation varies considerably among the published studies. While a smooth transition in biological communities is often conceptualized in association with estuarine fresh, brackish, and marine conditions, many studies have shown more distinct communities and the altogether absence of a brackish community. We explore these viewpoints in light of plant observations and soil and aquatic microbial analyses from permanent plots established on the Elbe River Estuary of northern Germany. Generally, two distinct communities were observed, a polyhaline assemblage towards the mouth of the system, and another that was associated with both the fresh and brackish mesohaline regimes further upriver. This was most pronounced among plant and soil bacteria communities, while aquatic 16S assemblages reflected little distinction at all. The proportion of marine classified taxa declined from the mouth to upriver and suggests that while the transition from marine to brackish or fresh vegetation falls within the sampled area, the same transition for microbial taxa could not be observed and may be further upriver. Thus, although we were able to identify two distinct communities, the “limit” of marine taxa was only evident for vegetation. While tidal and weather-related hydrology, as well as soil properties were also influential in distinguishing the communities, much of the variance remains unexplained. Further sampling, classification, and partitioning is necessary to determine the origin and/or autochthonous habitat, if any, for the Elbe River estuarine taxa.

**Geographic bounds:** bottom left: 53.556216°N, 8.824398°E

top right: 53.917760 °N, 10.155669 °E

## 1. Introduction

Estuaries are transitional landscapes composed of numerous gradients between land, freshwater, and sea. These gradients are thought to be fundamental to shaping the species assemblages from the broader regional biota to the more stress-tolerant and resilient taxa capable of surviving the range of physical and chemical characteristics typical of many estuarine systems (Teichert, et al., 2017; Engels & Jensen, 2010; Janousek & Folger, 2014; Attrill & Rundle, 2002). Although one classic and influential study suggests a smooth transition between marine and freshwater biota (Remane & Schlieper, 1972), subsequent studies have challenged this notion and instead suggest the presence of only marine and freshwater species, with no transitional biota otherwise referred to as “brackish” (Attrill & Rundle, 2002; Den Hartog, 1974; Telesh & Khlebovich, 2010; Whitfield, Elliott, Basset, Blaber, & West, 2012; Barnes, 1989). The semantics of estuarine salinity terms and the broad definition of “brackish waters” has also been noted and may contribute to the dispute (McLusky & Elliott, 2007; Elliott & McLusky, 2002; Davide, Marco, & Volpi, 2009). Thus, while numerous studies have confirmed the importance of salinity in shaping estuarine assemblages, there remains some debate on the presence of a true “brackish” community along a biological gradient. Further, other factors shaping these communities, such as flooding conditions, soil characteristics, and water physio-chemical conditions, among others, have received relatively little attention.

The Remane model for estuarine fauna (Remane & Schlieper, 1972) is based on findings from benthic fauna in the Baltic and presents the existence of three distinct biological communities within an estuary: fresh, brackish, and marine. Salinity is the primary driver of these differences, with the understanding that osmo-regulation abilities determine which community a species will belong to. While salinity tolerances and osmo-regulation are confirmed drivers of the differences in estuarine communities (Engels & Jensen, 2010; Tee, Waite, Lear, & Handley, 2021; Whitfield A. K., 2015), several studies have challenged the existence of true oligohaline biota and instead show evidence that this “brackish” zone of the estuary is instead inhabited by a mix of both marine and freshwater species (Attrill & Rundle, 2002). While this conclusion is based predominately on studies of macro-fauna and flora, microbial studies suggests both a brackish community that is simply a mix between fresh and marine communities, as well as truly autochthonous brackish taxa (Rocca, Simonin, Bernhardt, Washburne, & Wright, 2020; Herlemann, et al., 2011; Rieck, Herlemann, Klaus, & Grossart, 2015). Physiologically, another study concedes that while many brackish organisms are indeed of marine origin, they do express certain morpho-physiological traits that may constitute a brackish distinction (Cognetti & Maltagliat, 2000). Thus, there remains some disagreement on the true gradient nature of estuarine biota and the distinction of brackish community.

Other than salinity, additional drivers of biotic assemblages are confirmed (Wang, Chen, Zhang, Wang, & Kan, 2021). Estuarine hydrology is a result of the combined influence of precipitation-dependent surface waters, riverine freshwater runoff, groundwater, and tidally moving ocean waters (Snedden, Cable, & Kjerfve, 2013). Tidal dynamics often play a central role in controlling estuarine physio-chemical variability, but wind and fluvial forcings can also be relevant (Snedden G., 2006). Diurnal, seasonal, inter-annual, and even decadal variability in flooding regimes and freshwater availability have been attributed to changes in estuarine community assemblages (Janousek & Folger, 2014; Baptista, Martinho, Nyitrai, Pardal, & Dolbeth, 2015; Nodo, James, Childs, & Nakin, 2017). Soil and soil pore-water characteristics are also known to have some influence on marsh communities (Carling, et al., 2013), although at least one study documents little to no importance on microbial assemblages (Koretsky, et al., 2005).

This study aims to provide an initial assessment of the plant and microbial communities along the Elbe River Estuary in Northern Germany, focusing on the differences and similarities in assemblages along the estuarine salinity gradient, the partitioning of variability between the communities by different environmental variables, and an evaluation of the presence of true marine, brackish, and fresh communities. With the intention of developing a long-term research project, we aim to first establish an understanding of underlying physio-chemical patterns as they relate to biotic-interactions in the greater estuarine system.

## 2. Methodology

### 2.1 Study Site

The Elbe River Estuary encompasses approximately 170 river kilometers from the river’s mouth at the Wadden Sea in Northern Germany, to the extent of tidal influence at the Geesthacht weir east of Hamburg (Boehlich & Strotmann, 2008). Thirteen tributaries feed into the estuary as it widens into a funnel shape from 300-500 m wide at Geesthacht, to approximately 18 km wide at its mouth. The estuary holds economic importance as a major shipping route for international maritime traffic, especially to and from the port of Hamburg. Shipping channels and river embankments are regularly altered to maintain the necessary depths and flood controls for the harbors and surrounding populations. The active flood plain of the estuary, where tidal flooding affects estuarine vegetation, is limited to a few hundred meters between the mean high tide elevation and the constructed dikes, as well as on a few islands surrounded by the river. Along this strip, salt marshes occur at the estuary’s mouth, where salinity typically falls between 10-18 ppt (Engels & Jensen, 2010). Tidal freshwater and brackish marshes then occupy the floodplain for the remainder of the estuary, where salinity ranges between 0.5 and 10 ppt. Marsh communities typically follow an elevation zonation classified into three groups based on their flood frequency. Pioneer zone vegetation is flooded by almost each high-water event and thus twice a day, the low marsh is flooded during spring tides occurring with full and new moon twice a month, and the high marsh is flooded only during storm events, which are most frequent in autumn and winter months.

Although many studies on the Elbe and in other estuaries will refer to marine, brackish, and freshwater conditions as a general indicator of salinity, we will use an objective reference to describe site conditions and to avoid the confusion of semantics in this study. The Venice System for the Classification of Marine Waters According to Salinity was first proposed to avoid the ambiguity of “brackish” classifications (Battaglia, 1959). In this system, freshwater has a salt concentration less that 5 ppt, oligohaline concentrations are between 5 and 15 ppt, mesohaline is between 15 and 18 ppt and polyhaline is between 18 and 30 ppt. Although the system further describes higher salt concentrations, they are irrelevant to this study.

### 2.2 Study Design

We established nine permanent marsh measurement sites and six aquatic sites along 125 km of estuarine salinity and elevation gradients, from the mouth of the estuary to the city of Hamburg (Figure 1). The nine marsh sites were distributed among three locations spanning a 50 km stretch of river from the mouth to the village of Hohenhorst. At each of these locations, three sites were established along the elevation gradient: one at the pioneer zone, one at the low marsh, and one at the high marsh. At each marsh site, identical boardwalks were constructed to aid sampling and measurements and to limit soil and vegetation disturbance (Figure 2). These boardwalks provide access to three distinct zones within each site: a vegetation and soil sampling zone, a gas flux zone, and an undisturbed zone. In addition to the periodic sampling and measurements described in further sections, a series of autonomous and continuous measurements were also taken from an array of sensors placed into the soil and managed by a data logger at each site.

**Figure 1.**
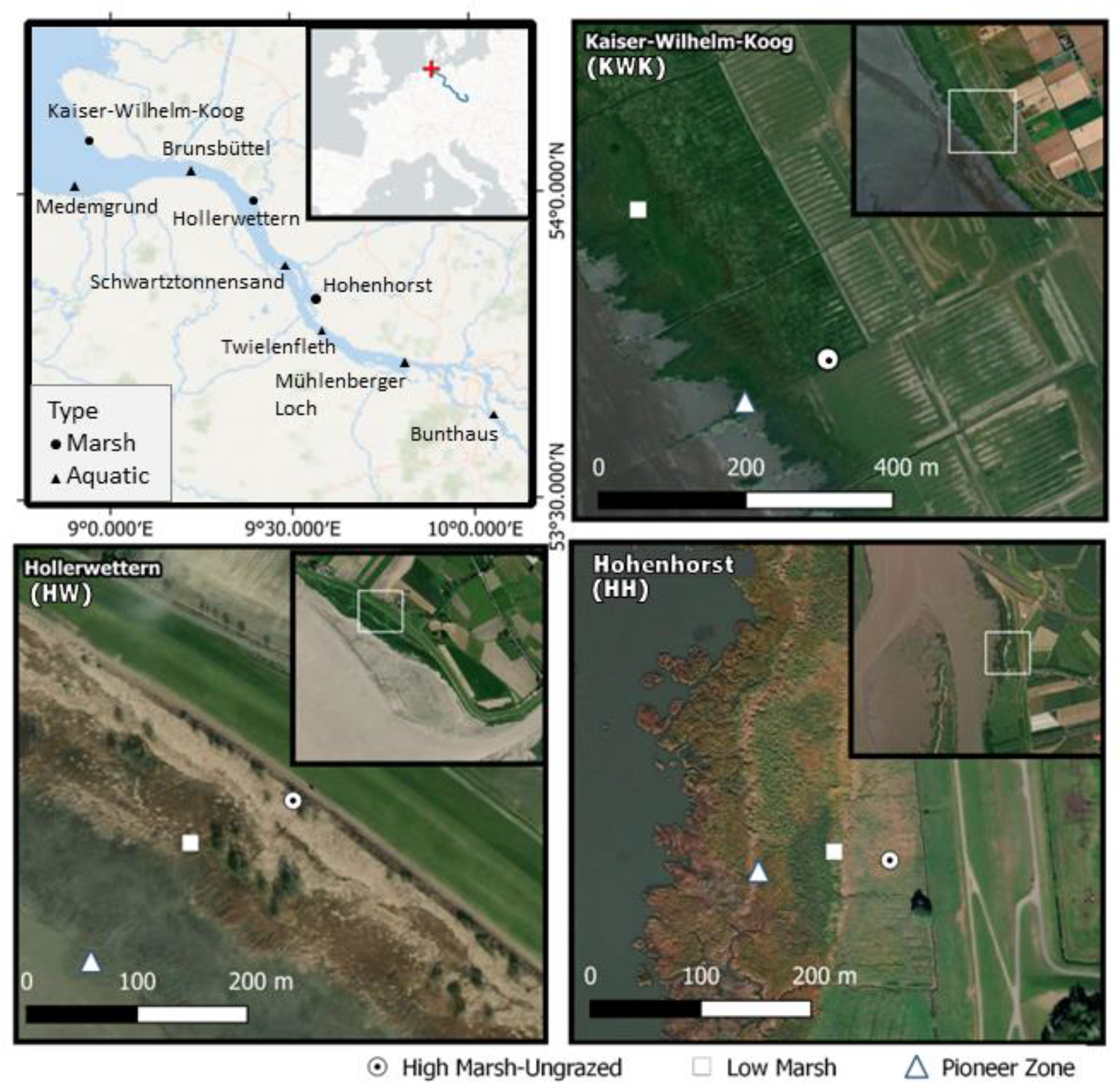
Nine marsh plots were established across three locations (Kaiser-Wilhelm-Koog, Hollerwettern, Hohenhorst) along with seven aquatic locations, where repeated sampling occurred along the Elbe River Estuary, Germany. Three sampling plots were demarcated at each of three marsh locations, one representing the pioneer zone, one the low marsh, and one in the high marsh. To allow for undisturbed sampling and measurements, boardwalks were constructed as depicted in Figure 2.

**Figure 2.**
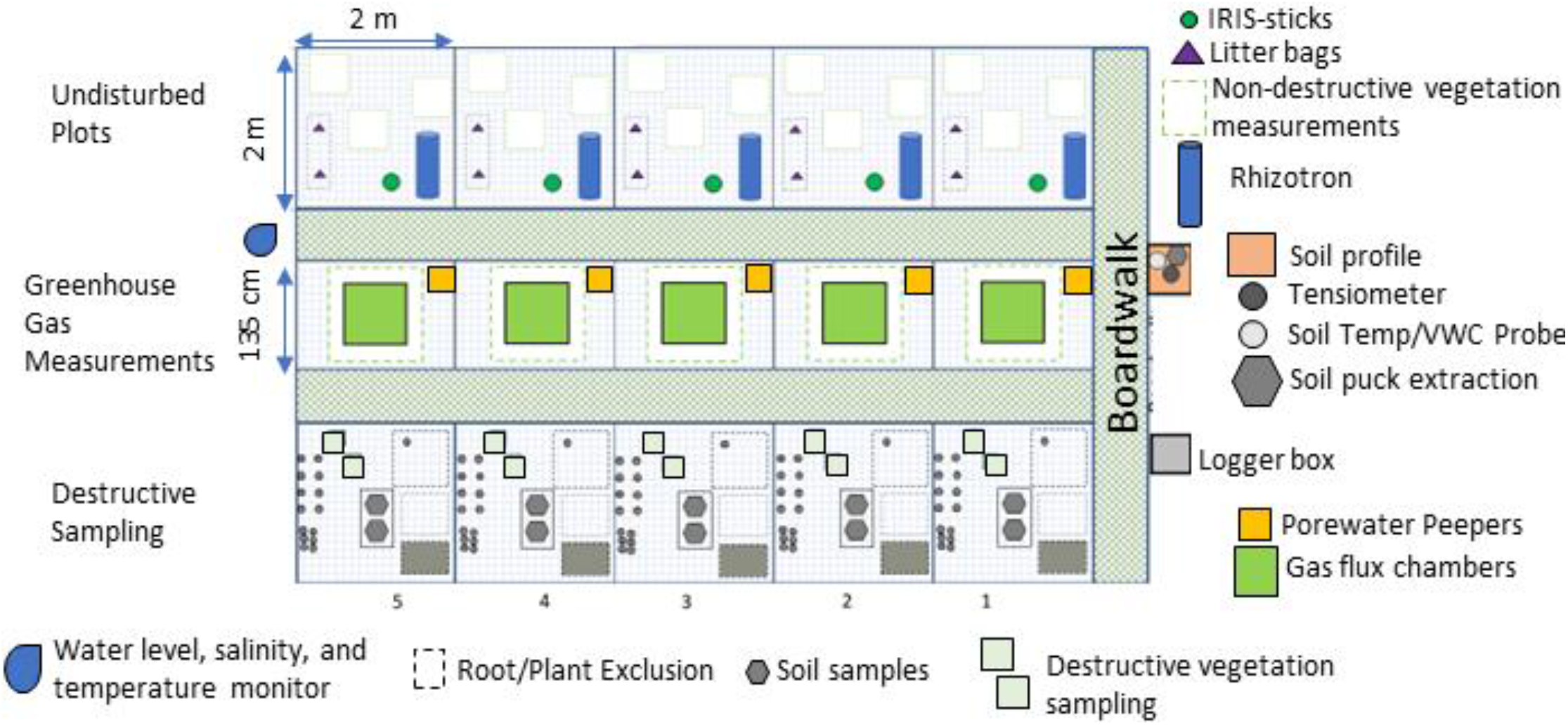
Marsh sampling and measurements were repeated at each study site among five replicate plots, which were located adjacent to boardwalks constructed to allow for easier access and less disturbance. Additional measurements beyond the scope of this study are also shown to demonstrate the project’s full scope.

### 2.3 Vegetation Surveys

Non-destructive species identification and coverage estimation was conducted in August 2022 in the undisturbed plots at each of the research sites over a two-day period. For vegetation surveying, five subplots (4m^2^ each) were selected along the 10 m boardwalk. Within each subplot, the coverage (%) of standing vegetation, bare soil and plant litter were assessed visually in two 60×60 cm quadrants (n=10). A coverage of 1% was assigned to plant species with a single occurrence, i.e. a single individual. The presence of seedlings was recorded but not included in further species analysis.

### 2.4 Soil Physio-Chemical Properties

Samples for soil characterization were collected in February and March of 2022 from each ungrazed marsh site. One soil sample was taken from all replicate plots (2 x 2 m, Figure 2) and separated into topsoil (0 – 10 cm) and subsoil (10 – 30 cm) samples. One aliquot of each sample was air-dried at room temperature and sieved to 2 mm for soil texture analyses by the sieving and sedimentation method for mineral soils (Müller, Dohrmann, Detlef, Rehder, & Eckelmann, 2009). The sand fraction (63 μm - 2000 μm) was analyzed by a vibratory sieve shaker (Retsch GmbH, Haan, Germany), while for the fine fraction (clay & silt) the Köhn-pipette fractionation with a Sedimat 4–12 (Umwelt-Geräte-Technik GmbH, Muencheberg, Germany) was used. Soil pH was measured in a 0.01 M CaCl_2_ solution (pH_CaCl2_) and in H_2_O (pH_H2O_) with a pH meter (MP230 GLP, Mettler-Toledo GmbH, Gießen, Germany). The electric conductivity (EC) was analyzed with a conductivity meter (WTW Cond 330i with TetraCon 325, Xylem Analytics Germany Sales GmbH & Co. KG, Weilheim, Germany). Another aliquot of each sample was dried at 105 °C until constant weight, sieved to 2 mm and ground for measuring inorganic and organic C contents with an elemental analyzer (soli TOC® cube, Elementar Analysensysteme GmbH, Germany) and total C and N contents (vario Max cube, Elementar Analysensysteme GmbH, Germany).

Soils at location were described and classified according to the IUSS Working Group WRB (2022) in November 2021, and in February and March 2022. A soil profile of minimally 100 x 100 x 60 cm was prepared for morphological descriptions and sampling. Various chemical analyses were performed on the soil samples to measure the soils’ ECse, N-NH_4_ and P-Olsen. ECse (electrical conductivity in saturation extract) is a proxy for readily soluble salts and is based on soil texture, organic carbon and electrical conductivity of a soil sample (WRB, 2022 chapter 8.4.28). The P-Olsen was determined by Olsen extraction method and photometrically measured.

### 2.5 Salinity and Flooding Dynamics

Salinity of groundwater and flooding dynamics were measured via installed piezometers at each marsh location, consisting of a roughly 6 cm diameter perforated PVC pipe with a CTD-Diver (VanEssen Instruments B.V.) pressure, temperature, and salinity sensor and data logger. Pipes were installed to a depth of roughly 1.5 meters, leaving 0.5 meters above the soil surface and the sensor hanging about 1.5 meters down from wire attached to the pipe’s cap. The exact length of these measurements varied by plot and was recorded upon installation for depth calculations. The sensor was programmed to capture measurements every fifteen minutes and data was collected routinely. Water depth was calculated by adjusting the recorded pressure to account for water density, which was calculated via water temperature and salinity, as well as barometric pressure, which was recorded at each location via a separate above-ground diver (VanEssen Instruments B.V.). The depth was then adjusted to reference the soil elevation by subtracting the depth of the sensor below the soil’s surface.

### 2.6 Soil Microbial Assays

#### 2.6.1 Soil sample collection

Soil samples were collected in September 2021 at each of the nine marsh sites. Soil cores were taken with a peat sampler (Eijkelkamp, Giesbeek, Netherlands) to collect samples at the surface and at 50 cm depth. Soil samples were stored at −80 °C until DNA extraction.

#### 2.6.2 DNA extraction from soil

0.5 g of soil was used for DNA extraction using the NucleoSpin Soil Kit (Macherey-Nagel, Düren, Germany), following the manufacturer’s protocol. Afterwards, the extracted DNA was analyzed in terms of yield and quality at a wavelength of 280 nm utilizing a Nanodrop spectrophotometer (NanoDrop 2000, Thermo Scientific, Waltham, USA).

#### 2.6.3 16S rDNA gene analyses

The sequencing and processing of the 16S rDNA variable regions V3-4 based on those described in Grüterich et al. (2024). Metabarcoding sequencing was executed at the Competence Centre for Genomic Analysis in Kiel, Germany. This process utilized the Illumina Nextera XT Index Kit, along with primers 341F (5’-CCTACGGGNGGCWGCAG-3’) and 785R (5’-GACTACHVGGGTATCTAATCC-3’), and the MiSeq Reagent Kit v3. Subsequent to demultiplexing, Cutadapt (v.4.4) (Martin, 2011) was employed for adaptor trimming of paired-end reads. The DADA2 pipeline (v.1.8) was used for read filtering and ASV inference, with specific criteria (maxEE=2, maxN=0, truncQ=2, truncLen=260,210). The taxonomic assignment of merged and corrected reads was accomplished using the SILVA database (v.138.1).

### 2.7 Aquatic Microbial Assays

#### 2.7.1 Aquatic sample collection

Aquatic samples were collected in May and July 2021, and February, May, June, November 2022 at each of the 6 aquatic stations. Samples were taken with horizontal sampler as described in Tobias-Hünefeldt (2024). Two-hundred and fifty mL of water was filtered through 5 μm and 0.22 μm Durapore filters (Merck: SVWG04700, GVWP04700). Filters were stored at −20 °C until DNA extraction.

#### 2.6.2 DNA extraction and sequencing from aquatic filters

DNA was extracted as in Nercessian et al. (2005). Metagenomic sequencing was performed at Ramaciotti Centre for Genomics (Sydney, Australia) and the Competence Centre for Genomic Analysis Kiel (Kiel, Germany). Samples were prepared for sequencing with the Illumina DNA prep kit, and sequenced on a NovaSeq 6000 platform (Illumina, San Diego, CA, USA). Raw metagenomic sequenced underwent amplicon extraction using mTAGs, as described in (Salazar, Hans-Joachim, Hildebrand, Acinas, & Sunagawa, 2022).

### 2.8 Weather Data

Weather stations were installed at each of the three marsh locations. Instruments include the SN500SS Apogee four-component net radiometer to track both short- and long-wavelength radiation and down-welling from the atmosphere and emitted from the ground. Also included is a WindSonic4 Gill ultrasonic anemometer, installed 2 meters above the surface, measuring wind speed and direction. The HygroVUE10 Temperature and Relative Humidity Sensor records conditions at the 2-meter mark above the surface. Finally, the unheated Campbell Scientific tipping bucket rain gauge measures rainfall. All climate variables are measured continuously with a temporal resolution of 5 minutes.

### 2.9 Data Analyses

Data processing and statistical analyses were done in R (R Core Team, 2023). Community assemblages were primarily evaluated through the vegan package (Oksanen, et al., 2022). Differences in community assemblages were evaluated through PERMANOVA by way of the *adonis2*() function. The same was also done as pairwise evaluations for each pair of communities via the *pairwise.adonis*() function from the pairwiseAdonis package (Martinez, 2020). Ordinations of the communities via non-metric multidimensional scaling (NMDS) was accomplished through the *metaMDS*() function, and environmental variables were fit to these ordinations via the *envfit*() function, both from vegan. Environmental variables for the ordination were first selected for through variance partitioning via the *varpart*() function from vegan. We used this function to find the optimal combination of environmental variables that were not correlated with each other and that maximized the explained variability in the ordination.

Classification of taxa into “marine”, “brackish”, and “freshwater” groups was accomplished via queries to the World Register of Marine Species (WoRMS) (Worms Editorial Board, 2024). We used the worms package in R (Chamberlain, 2023) to query taxa names and combined the “isMarine”, “isBrackish”, and “isFreshwater” variables to create groups depending on if only one was indicated (Marine, Brackish, or Freshwater) or if some combination of the three was indicated (Brackish & Marine, Brackish, Marine, & Freshwater, etc.).

### 2.10 Data availability

Raw reads of the 16S rDNA analysis were deposited at the European nucleotide archive ENA under the project accession number PRJEB54081 (Biosamples: SAMEA110291994 - SAMEA110292011 and SAMEA113954754 - SAMEA113954807). ASV tables for the 16S rDNA analyses were deposited at the ZFDM repository of the University of Hamburg (https://doi.org/10.25592/uhhfdm.13933).

All other data will be made available on the project data repository (https://www.fdr.uni-hamburg.de/communities/rtg2530), in some cases after an embargo period of 24 months has passed since the termination of individual studies.

## 3 Results

### 3.1 Salinity and Tidal Dynamics

Salinity measurements at the sampling locations ranged from 0.1 to 27.6 ppt, although mean values at each location ranged from 0.5 to 17 ppt and fell mostly within either mesohaline or oligohaline conditions (Figure 3). Some sites fell on the extremes of these conditions. Bunthaus, with mean salinities of about 0.45 fell on the maximum limit of a freshwater classification although many measurements were oligohaline. Conversely, Kaiser-Wilhelm-Koog had a mean salinity of 16.8, but many readings exceeded 18 ppt and thus classify as polyhaline. Generally, salinity standard deviations averaged about 37% of the mean values and ranged from 11% to 54%. Daily flood frequency among the marsh sites ranged from about once every one-hundred days to twice a day and on average increased upriver from Kaiser-Wilhelm-Koog to Hohenhorst and from high marsh to the pioneer zone. At the seaward site of Kaiser-Wilhelm-Koog, flood frequency increased from once every ten days at the high marsh to once every 1.4 days at the pioneer zone. At the mid-estuarine site of Hollerwettern, these values were every one-hundred days at the high marsh to every 12 hours at the pioneer zone, and at Hohenhorst, they were every three days at the high marsh and every 20-22 hours at the low marsh and pioneer zones.

**Figure 3.**
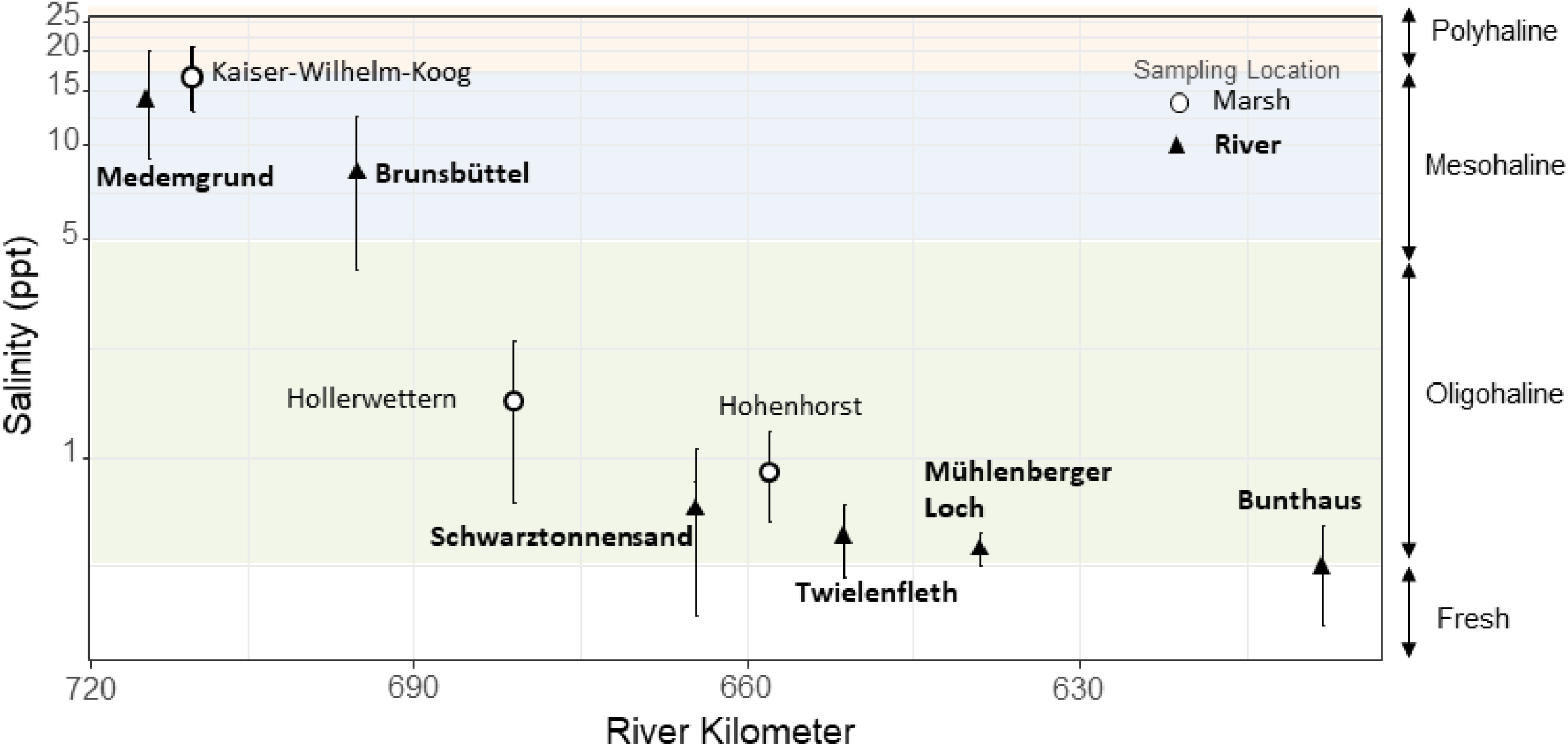
Salinity means and standard deviations measured at the sampling locations as represented by their distance along the Elbe River. Note the logarithmic scale of the salinity axis, which demonstrates the variability in salinity at each site, but otherwise misrepresents differences in mean values between the sites. Two aquatic sites fall on the salinity boundary between fresh and oligohaline definitions, and one marsh site falls on the boundary between mesohaline and polyhaline conditions.

Most flooding events across the sites lasted between 3 and 10 hours, with the lowest median flood length of 2.8 hours occurring at the mid estuarine (Hollerwettern) high marsh and the longest median flood length of 32 hours occurring at the upper estuarine (Hogenhorst) high marsh. Conversely, most of the non-flooded periods lasted between 4 and 58 hours (about 2 and a half days), but the longest median non-flooded length of 978 hours (about 1 and a half months) occurred at the mid estuarine high marsh.

### 3.2 Microclimatic characteristics

The microclimatic conditions at the three marsh locations are relatively similar, with the exception of rainfall and short-wave radiation. Average annual near-surface air temperature at the three sites is 10.4° C to 10.6°C, which is consistent with the mean air temperature of 9.3° C for the Hamburg region, based on a global reanalysis climate data set spanning the period 1979 to 2019 (Hersbach, et al., 2020). Thereby, the average air temperature for the years 1979 to 1983 compared to the years 2015 to 2019 increases from 8.6° C to 10.3° C, which illustrates the warming trend over the last decades. Within a year, the measured average daily air temperature ranges from –4.8° C to 26.7° C. The annual rainfall at the sites amounts to 668 mm/a, ranging from 593 mm/a at Hollerwettern to 737 mm/a at Kaiser-Wilhem-Koog. This is slightly less than the 1979 to 2019 average of 782 mm/a for the Hamburg region based on ERA 5 data. In contrast to air temperature, a long-term trend in rainfall amount is not obvious here. The majority of total rainfall occurs in the winter months, 217 mm from December to February compared to 125 mm from June to August. Daily average down-welling short-wave solar radiation ranges from 26 W/m^2^ in the winter months to 193 W/m^2^ in the summer months. Thereby, the station at Hollerwettern received considerably less radiation, 108 W/m2, in summer than the other two stations. Down-welling long-wave radiation (near infrared) is more evenly distributed throughout the year and between stations varying from 308 W/m^2^ in winter to 337 W/m^2^ in summer. Relative humidity of the air is constantly high at the marsh locations with lowest average values of 76% in the summer months and 92% in winter. Only 11 to 14 days in a year show average relative humidity below 60%. The wind speed shows a weak seasonal pattern with higher values in autumn and winter, and median values of 2.3 m/s to 2.7 m/s over the year. Two periods in March, however, show storm conditions with wind speed reaching peak values of up to 58 m/s.

### 3.3 Soils

Different soil reference groups have been assigned to the estuarine soils; Solonchaks and Gleysols, reflecting the different dominant soil forming processes characterizing the soils. The soils of the estuarine mouth at Kaiser-Wilhelm-Koog (KWK) are characterized by high concentrations of soluble salts reflecting the strong influence from the marine dominated tidal floods. Whereas, the mid-estuarine (Hollerwettern, HW) and upriver (Hohenhorst, HH) marsh soils are characterized by iron and aluminum chemistry reflecting the strong influence of ground water and riverine tidal floods. Averaged electrical conductivity (EC) at the high, low, and pioneer zone marshes were 28.9, 10.8 and 24.0 dS m^−1^, respectively, at KWK and 1.7, 1.6 and 1.1 dS m^−1^ at HH. The same values, minus the grazed high marsh, for Hollerwettern were 1.6, 5.3, and 2.5 dS m^−1^.

Mineral nitrogen levels (N-NH4) ranged from 3.8 to 7.9, 0.8 to 4.6 and 2.2 to 147.7 μg N g-1 dry weight at KWK, HW, and HH marsh respectively. In the same order, Phosphorus levels were 23 to 46, 3.6 to 30.6 and 9.2 to 33.3 mg P(-Olsen) per kg soil, respectively.

Soil texture across locations and elevations were dominated by sandy or silt loam. Silt averaged 50 % content, followed by sand at 31 % and clay at 19%. There were no significant differences (wilcox-test) in the percent content of any component between the different elevations when pooled across the locations. However, there were significant differences in contents at each of the individual locations. The most upriver site (Hohenhorst) had generally higher clay content compared to the other sites, while the mid-estuarine site (Hollerwettern) was partly dominated by sand contents >60%. At the polyhaline site (Kaiser-Wilhelm-Koog), silt content decreased significantly from 68% in the high marsh to 48% in the pioneer zone, while sand content increased along the same gradient from 8% to 33%. Clay content was significantly higher in the high marsh at 23% than at both the low marsh and pioneer zone at 15%. Total organic carbon (TOC) decreased significantly from 3.3% in the high marsh to 2.2% in the pioneer zone, with all locations pooled together. This general pattern held for the Kaiser-Willelm-Koog and Hollerwettern (oligohaline) location, but not the Hohenhorst location, which held a statistically similar TOC percentage with a mean of 4.2% across elevations. This was significantly higher than the mean percentage at both of the other locations.

The soil redox index increased consistently from the mouth of the estuary at Kaiser-Wilhelm-Koog to the uppermost location at Hohenhorst, and from the high marsh to pioneer zone sites at each location. On average, the index measured 0.23, 0.45 and 0.45 at the three locations moving upriver, respectively. There was a general trend of increasing redox index with decreasing electrical conductivity, however the high marsh sites at the two oligohaline locations (HW and HH) held relatively low redox indices for their respective conductivities.

### 3.4 Vegetation

Forty-four plant species were identified within the survey plots of the estuary. These species belonged to twenty-three distinct families, with nearly 23% of the species belonging to the Poaceae family and the remaining families harboring only one or two of the identified species. The Kaiser-Wilhelm-Koog, Hollerwettern, and Hohenhorst sites held 18, 19, and 14 unique species, respectively, across their plots. Individual plots from the polyhaline/mesohaline location (KWK), however, held significantly more species (mean = 10) than both the oligohaline sites at Hollerwettern (mean = 7.7, mean difference = 1.4, p<0.01) and Hohenhorst (mean = 5.7, mean difference = 2.2, p<0.001), both of which held statistically similar species numbers to each other (p>0.5).

### 3.5 Soil Microbial Communities

We detected 719 unique microbial genera from 282 families and 30 different phyla among the soils sampled. Within this dataset, also archaeal sequences are included, but only represented with 9 individual genera and a maximum abundance of 1.3% within all samples. It is possible that the bias in archaeal sequence abundance stems from the challenge of effectively lysing their tough cell walls using conventional DNA extraction methods, leading to underrepresentation. The median relative abundance for the most abundant phylum in each sample was 5% and ranged from 2% to 42%. *Campylobacterota* was the most abundant phylum in 40% of the samples with a mean relative abundance of 16% across all samples. This was due almost entirely to the abundance of *Sulfuricurvum*, which was the most abundant genus in 30% of the samples, with a median and max abundance of 19% and 42%, respectively. The next most abundant genera were an MND1 group, and an unknown group referred here as “subgroup 1”, which were the most abundant groups in 10% and 7% of the samples, respectively. The remaining 53% of the samples were dominated by fifteen other genera.

The soils of the poly/mesohaline plots at Kaiser-Wilhelm-Koog and those of the mid-estuarine oligohaline plots at Hollerwettern held statistically more taxonomic genera than the upriver oligohaline soils at Hohenhorst (mean difference = 19 genera, p<0.05). There was no significant difference in soil microbial diversity between KWK and HW (mean difference =7 genera, p < 0.5).

### 3.6 Aquatic Microbial Communities

4760 distinct 16S genera were identified in the aquatic samples, although the most abundant of these were unidentified. The most and second most abundant genera in each of the 180 samples were unknown taxa. The top-ranked known genus was identified as “hgcI clade”, which was the third most abundant taxon in 10% of all samples, followed by *Flavobacterium* in about 2% of the samples and *Amoebophyra, Woeseia*, and MB11C04 each were the third ranked taxon in one sample.

There were no significant differences in the number of unique genera between the sampling points. However, Brunsbüttel at the mouth of the estuary held on average 343 more genera than Bunthaus at the upriver end of the sampling transect, a difference that was nearly significant (ANOVA, p < 0.06).

### 3.7 Community Partitioning

Distinct communities within the different salinity regimes were detected for both vegetation and soil 16S microbial assemblages, and to a lesser extent in the aquatic 16S microbes (Figure 4). For vegetation, all three collective plant communities at each location were distinct and significantly different than the others (Pairwise ADONIS: p<0.001, R^2^=0.3), as were those representing the oligohaline (HW and HH) and the polyhaline/mesohaline (KWK) salinity regimes (Pairwise ADONIS: p<0.001, R^2^=0.2). Within location, all communities at each of the three elevation groups were also distinct and significantly different from the others (Pairwise ADONIS: p_KWK < 0.005, R^2^_KWK=0.6; p_HW < 0.005, R^2^_HW = 0.6; p_HW < 0.005, R^2^_HW = 0.3).

**Figure 4.**
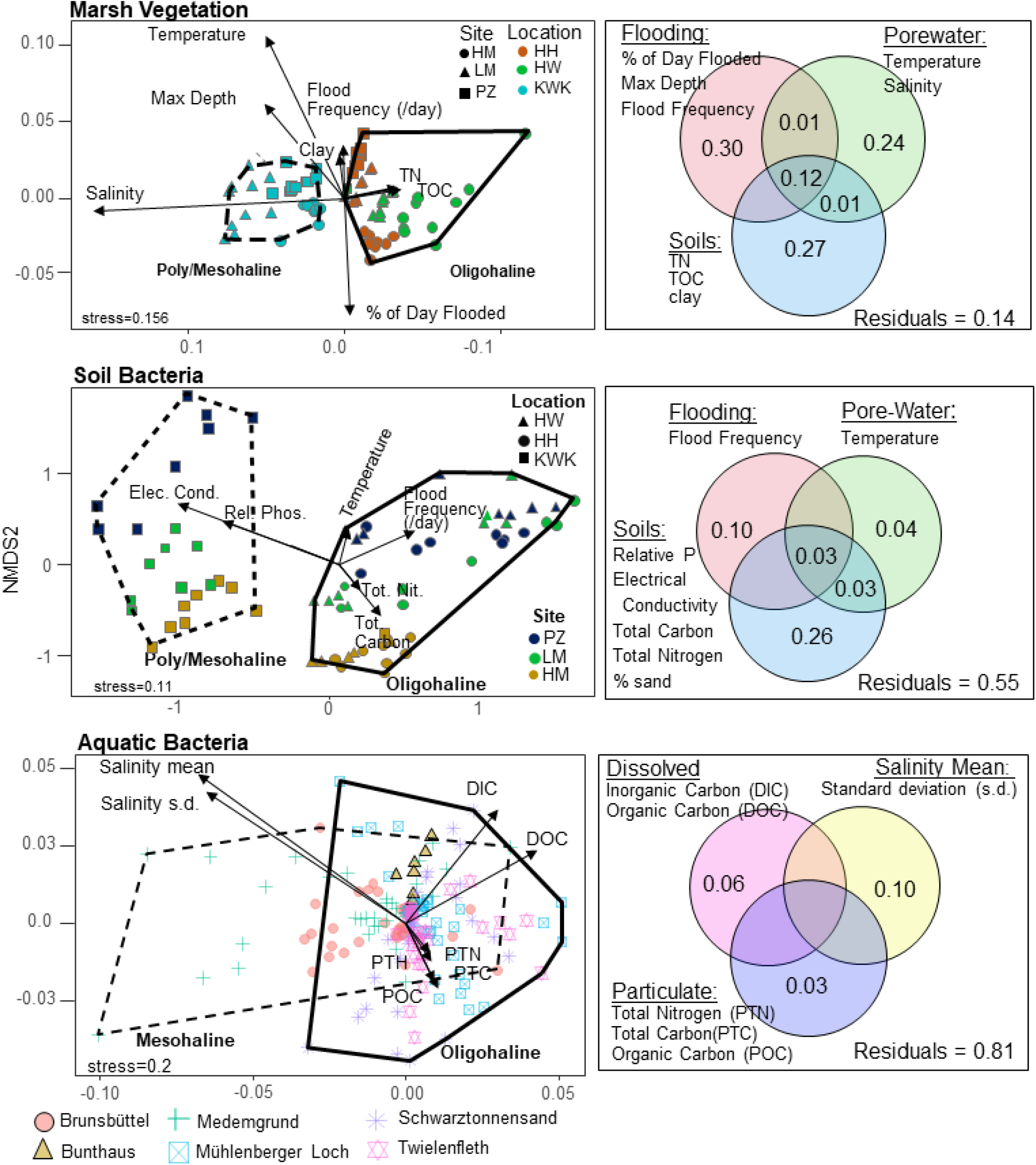
Nonmetric multidimensional scaling ordinations of the Elbe River estuary plant, soil 16S microbial, and aquatic 16S microbial communities (left panels), with their respective variance partitioning of environmental variables (right panels). At least two distinct communities, one oligohaline and one poly/mesohaline can be identified in the vegetation and soil microbial samples, with a much weaker distinction detected in the aquatic microbial samples.

Soil 16S microbial communities were also distinct among the two salinity regimes (Pairwise ADONIS: p<0.001, R^2^=0.2) but the communities at the two oligohaline locations, Hohenhorst and Hollerwettern, were statistically similar and non-distinct (Pairwise ADONIS: p>0.5, R^2^=0.04). Between elevations at each location, soil 16S communities were not always distinct, with the pioneer zone and low marsh communities being non-distinct at the oligohaline locations (Pairwise ADONIS: p_HW > 0.1, R^2^_HW = 0.1; p_HH > 0.5, R^2^_HH = 0.1), but not at the poly/mesohaline location (Pairwise ADONIS: p_KWK < 0.05, R^2^_KWK = 0.4). Depth was also an important factor in differentiating microbial soil communities. When pooled together, the surficial and depth samples from all sites were statistically different, albeit relatively weakly (p=0.001, R2=0.1). This difference was statistically significant and stronger, however, when tested individually at each study site (minimum R^2^ = 0.46, max p = 0.04).

Significant differences in 16S communities were also found between all but two of the aquatic salinity regimes, although these differences were relatively weak in comparison with the soil and vegetation communities (Pairwise ADONIS: p < 0.05, R^2^ > 0.04). The two exceptions were between the polyhaline/mesohaline community at Medemgrund and the collective mesohaline communities upriver (Pairwise ADONIS: p < 0.3, R^2^ = 0.04).

Generally, variance partitioning in the distinction of the communities was best explained in the vegetation assemblages (86% explained variance) and less so in the soil 16S (45% explained variance) and aquatic 16S (19% explained variance) assemblages. Flooding, porewater, and soil variables each explained about 30% of the vegetation community variability. In soil 16S communities, most of the variability was explained by soil characteristics (26%), while flooding and porewater explained considerably less (10% and 4%, respectively). In aquatic 16S communities, the standard deviation of salinity was responsible for most of the explained variability (10%), while dissolved and particulate carbon and nitrogen contents together explained the remaining 9% of the explained variability.

### 3.8 Marine classification

Eleven (25%) of the vegetation species were identified and classified by WoRMS (Figure 5). The only classified marine plant species was *Atriplex littoralis*, which was found only at the mouth of the estuary in KWK. The remaining classified plant species were brackish and brackish-fresh. In soil bacteria, referencing the WoRMS database returned 83 genera (6%) that were identified and classified. Of these, 76 were classified as marine, four as mixed, and three as brackish & marine. For aquatic bacteria, two-hundred and eighty-three of the genera (6%) were identified and classified in the WoRMS database. Two-hundred and fifty-one of these were classified as marine, 4 as brackish, one as brackish-marine, and 17 as mixed. Along the salinity gradient, an expected decline in marine classified species from the estuary’s mouth to upriver occurred in all groups. For vegetation, the transition from some marine influence to no marine influence occurred between the estuary’s mouth at KWK and the mid-estuary site at Hollerwettern. For bacteria, although there was a decline moving upriver, all samples included marine classified taxa.

**Figure 5.**
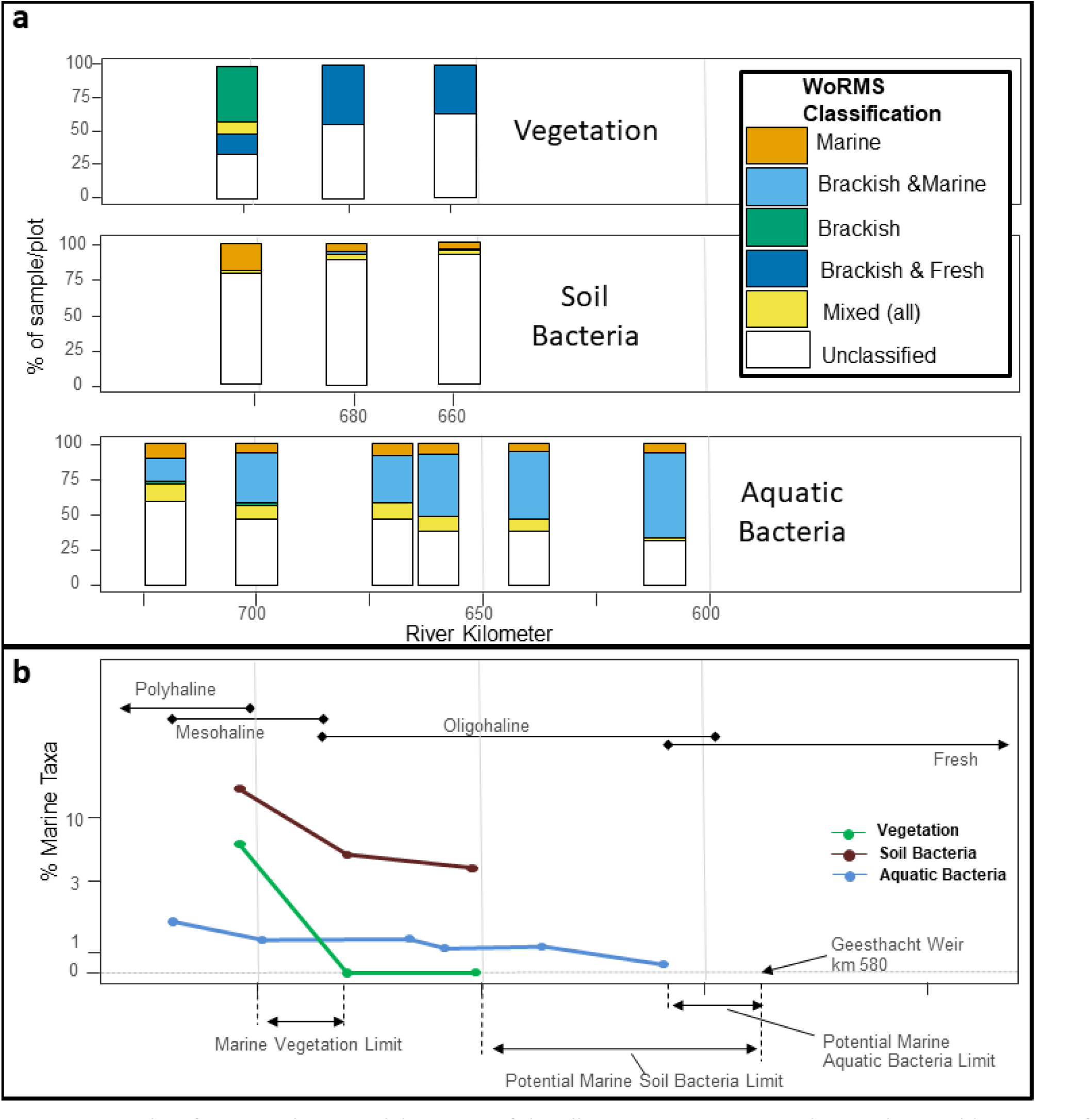
Taxa classifications along 125 kilometers of the Elbe River Estuary, according to the World Register of Marine Species (WoRMS). Panel a shows all classifications for each group at each sampling location. Panel b shows only the marine proportion as a function of the river distance. Although many taxa were unclassified in the database, among those that were, the proportion of marine species decreases moving upriver. For vegetation, no marine species were recorded before kilometer 700, suggesting the marine transition lies between kilometer 680 and 700 (panel b). For bacteria, marine species were recorded at all stations, suggesting their limit is at an unknown distance upriver. Our indication of upriver proportions for bacteria (shaded areas, panel b), assume they do not cross the Geesthacht Weir.

## 4 Discussion

Estuaries are largely conceptualized and modeled as gradients in physical and chemical conditions between land and water, fresh and marine. Biological organization within these systems has also been shown to follow a similar pattern, but recent studies have challenged this concept. These studies have suggested that instead of a smooth transition in community assemblages between marine, brackish, and freshwater conditions, there exists only two true species pools corresponding to marine and freshwater environments. Everything in between these two salinity regimes is a mix of the two principal communities, with marine species dominating most of the brackish conditions. Additionally, while salinity has been singled out as the primary driver of these differences in community assemblages, flooding conditions and soil characteristics have received relatively little attention as potential drivers. Our results indicate that while there was generally a smooth transition in the proportion of marine classified taxa along the estuary’s length, the nature of this transition was different for vegetation, soil bacteria, and aquatic bacteria. In community analyses, we also found differences in the distinction between assemblages depending on the biota of interest. Thus, our preliminary observations from vegetation and microbial communities on the Elbe River estuary do not provide unequivocable evidence for either the gradient or marine-fresh models of estuarine biotic assemblages, although traits of both models are evident.

For vegetation and soil bacteria, there was clear evidence for only two primary communities, one oligohaline and one meso/polyhaline. However, the contribution of marine taxa to these communities was not consistent and did not support a single model of marine influence on estuarine communities. In vegetation, only one species was classified as marine and only at the mouth of the estuary, whereas in soil bacteria, the contribution of marine taxa decreased gradually moving upriver and all samples included some marine taxa. Further, while a similar trend was observed in aquatic bacteria, there was no strong distinction in communities as was observed in soil bacteria. These differences may be due to the previously acknowledged inherent variability in estuaries, with no single model applicable to all (Whitfield, Elliott, Basset, Blaber, & West, 2012). Taxa mobility likely also contributes to this (Matela & Obolewski, 2022), where vegetation and soil bacteria are less mobile than aquatic bacteria and thus less capable of avoiding high salinity stress. This is further supported by a complete lack of any freshwater classified taxa, a pattern that has been noted previously in other systems (Whitfield A. K., 2015), and that likely results from mobile organisms avoiding saline environments and non-mobile organisms unable to survive. However, limitations in WoRMS classifications as well as in our sampling spatial and temporal resolution may also contribute to some of these deviations from proposed models. Our project, for example, did not sample any truly freshwater or marine conditions and the WoRMS database is likely biased towards marine taxa, rather than freshwater taxa.

For both vegetation and microbes, diversity in the present study was highest at the poly/mesohaline sites. Several studies have shown the opposite for both microbial (Campbell & Kirchman, 2013; Osterholz, Kirchman, Niggemann, & Dittmar, 2018) and plant (Watson & Byrne, 2009; Janousek & Folger, 2014) diversity in other estuaries and suggest that the stress of elevated salinity constrains the potential species’ pools, especially in intermediate salinity zones (greater than five and less than 30 PSU). Attrill (2002) suggested that salinity variability, rather than absolute mean salinity, was the greatest contributor to estuarine organismal distribution and the greatest constraint on diversity. Our findings also do not support this, as the most saline sites held both the highest variability in salinity and the highest diversity of both vegetation and microbes. Still, other studies suggest that salinity is not the primary driver of estuarine species assemblages and that other factors, such as elevation, hydrology, or edaphic conditions are most influential (Boey, Mortimer, Couturier, Worrallo, & Handley, 2022).

While salinity underlies many of our assumptions regarding the observed patterns in Elbe estuarine biology, soil and hydrology also played some role in partitioning communities. As represented by the vectors of environmental variables in our ordination (Figure 4), pore-water salinity and soil electrical conductivity (also a proxy for ion content) were strongly correlated (as denoted by their vector lengths) with the direction of separation between the distinct communities. This is most notable in the vegetation and soil bacteria communities, but less so in the aquatic bacteria. Also of note, however, is the importance of flooding dynamics in distinguishing communities along the marsh elevation gradient, which is oriented along the vertical axis (NMDS2) in Figure 4. Again, this is most notable for vegetation and soil bacteria communities, a pattern that has been observed previously (Engels & Jensen, 2010; Janousek & Folger, 2014; Griffin, et al., 2020). As for aquatic bacteria, the suite of available environmental variables was least explanatory in comparison to vegetation and soil bacteria, although salinity explained the largest portion of the variance. Further sampling and testing against novel environmental variables, as well as more complex biotic interactions, may help elucidate the variability in aquatic bacteria assemblages.

Generally, our results both agree with and contradict several previous studies that demonstrate estuarine community assemblages. While a smooth transition in marine taxa presence does generally follow the classic Remane model (Remane & Schlieper, 1972), the transition was not consistent among the sampled communities, as suggested by Whitfield et al (2012). Further, while distinct communities were observed along the estuary (Attrill & Rundle, 2002), the distinction was also not consistent among the vegetation, soil bacteria, and aquatic bacteria communities. Also in contradiction to Attrill & Rundle (2002), who proposed brackish communities are a mix of marine and freshwater biota, we found no strictly freshwater classified taxa in our samples and instead found a mix of all classifications, again varying depending on the community in question. Thus, while it’s clear that patterns in community transitions within estuaries depends largely on the taxonomy (e.g. tracheophytes vs. prokaryotes) and nature (e.g. terrestrial vs. aquatic, sessile vs. mobile), much of these disagreements may be explained by a misuse or misunderstanding of the term “brackish” among the studies and including within the WoRMS database. We therefore caution against this term and instead encourage use of defined salinity regimes as represented by the Venice System (Battaglia, 1959). We also encourage future sampling to include extreme ends of the salinity gradient, which may help elucidate the origin and distribution of distinct communities within the Elbe River Estuary.

## 5 Author Contributions

**Benjamin Branoff**: conceptualization, methodology, formal analysis, data curation, writing – original draft, writing – review and editing, visualization. **Luise Grüterich**: formal analysis, investigation, data curation, writing – review & editing. **Monica Wilson**: formal analysis, investigation, writing – review & editing. **Sven P. Tobias-Hunefeldt**: formal analysis, investigation, data curation, writing – review & editing. **Youssef Saadaoui**: formal analysis, investigation, data curation, writing – review & editing. **Julian Mittmann-Goetsch**: formal analysis, investigation, data curation, writing – review & editing. **Friederike Neiske**: formal analysis, investigation, data curation, writing – review & editing. **Fay Lexmond**: formal analysis, investigation, data curation, writing – review & editing. **Joscha Becker**: formal analysis, investigation, data curation, writing – review & editing. **Hans-Peter Grossart**: resources, writing – review & editing, supervision, project administration, funding acquisition. **Philipp Porada**: resources, writing – review & editing, supervision, project administration, funding acquisition. **Wolfgang R. Streit**: resources, writing – review & editing, supervision, project administration, funding acquisition. **Annette Eschenbach**: resources, writing – review & editing, supervision, project administration, funding acquisition. **Lars Kutzbach**: resources, writing – review & editing, supervision, project administration, funding acquisition. **Kai Jensen**: resources, writing – review & editing, supervision, project administration, funding acquisition.

## Notes

https://www.fdr.uni-hamburg.de/communities/rtg2530

### Competing Interest Statement

The authors have declared no competing interest.

